# Mosquito immune cells enhance dengue and Zika virus dissemination in *Aedes aegypti*

**DOI:** 10.1101/2024.04.03.587950

**Authors:** David R. Hall, Rebecca M. Johnson, Hyeogsun Kwon, Zannatul Ferdous, S. Viridiana Laredo-Tiscareño, Bradley J. Blitvich, Doug E. Brackney, Ryan C. Smith

**Affiliations:** Interdepartmental Program in Genetics and Genomics, Iowa State University, Ames, Iowa; Department of Plant Pathology, Entomology and Microbiology, Iowa State University, Ames, Iowa; Center for Vector-Borne and Zoonotic Diseases, Department of Entomology, The Connecticut Agricultural Experiment Station, New Haven, Connecticut; Department of Veterinary Microbiology and Preventative Medicine, Iowa State University, Ames, Iowa

**Author notes:** These authors contributed equally and are listed alphabetically (David R. Hall, Rebecca M. Johnson, Hyeogsun Kwon).

## Abstract

Mosquito-borne viruses cause more than 400 million annual infections and place over half of the world’s population at risk. Despite this importance, the mechanisms by which arboviruses infect the mosquito host and disseminate to tissues required for transmission are not well understood. Here, we provide evidence that mosquito immune cells, known as hemocytes, play an integral role in the dissemination of dengue virus (DENV) and Zika virus (ZIKV) in the mosquito *Aedes aegypti*. We establish that phagocytic hemocytes are a focal point for virus infection and demonstrate that these immune cell populations facilitate virus dissemination to the ovaries and salivary glands. Additional transfer experiments confirm that virus-infected hemocytes confer a virus infection to non-infected mosquitoes more efficiently than free virus in acellular hemolymph, revealing that hemocytes are an important tropism to enhance virus dissemination in the mosquito host. These data support a “trojan horse” model of virus dissemination where infected hemocytes transport virus through the hemolymph to deliver virus to mosquito tissues required for transmission and parallels vertebrate systems where immune cell populations promote virus dissemination to secondary sites of infection. In summary, this study significantly advances our understanding of virus infection dynamics in mosquitoes and highlights conserved roles of immune cells in virus dissemination across vertebrate and invertebrate systems.

## Introduction

In recent decades, climate change and globalization have driven the expansion of *Aedes albopictus* and *Aedes aegypti* from their native tropical and subtropical habitats to more temperate environments across the globe, such that they are now present on every continent except Antarctica^1–3^. As the primary vectors of dengue (DENV), Zika (ZIKV), and many other arboviral diseases, the expansion of these mosquito species has increased the incidence of DENV by 30-fold in the last 50 years, reaching an estimated 100 million clinical cases per year^4^. Likewise, arboviral outbreaks, such as the emergence of ZIKV in the Americas in 2015 and 2016, have underlined the increasing risk and public health threat presented by these mosquito species and the viruses they transmit^5^. With the risk of arbovirus transmission likely to continue to increase and further expand into new areas in the future^6–8^, understanding the key factors that influence arbovirus transmission by the mosquito host is of significant importance.

After entering the mosquito through an infectious blood meal, viruses must overcome multiple tissue barriers in the mosquito host in order to reach the salivary glands and be transmitted to a new host^9–11^. While virus uptake and replication in the midgut are an essential first step to mosquito infection, the manner by which the virus disseminates from the midgut to other mosquito tissues remains poorly understood. Several potential routes of virus escape into the hemolymph and secondary tissues have been suggested, including routes through the tracheal system^12,13^, damage to the midgut basal lamina^14–17^, and infection of the visceral muscles^18^. Although each of these potential routes, or combinations thereof, may contribute to virus dissemination, recent evidence suggests that the midgut basal lamina may be the most significant barrier. Several studies have shown that damage to the basal lamina resulting from an additional blood meal significantly enhances virus dissemination, reducing the extrinsic incubation period^14,17,19^.

Mosquito immune cells, known as hemocytes, serve as important immune sentinels that circulate in the hemolymph and have integral roles in pathogen recognition, immune signaling, and wound healing^20,21^. Hemocytes are found either in circulation or attached to different mosquito tissues as sessile cells^22^ and can readily become infected by virus^23–27^. As a result, it has been suggested that hemocytes may serve as an additional tropism for virus replication in the hemolymph^25^. While evidence supports that mosquito and *Drosophila* hemocytes contribute to antiviral defenses^26,28–31^, the potential that virus-infected hemocytes may also promote virus dissemination in the mosquito host has not been previously explored.

While previous studies in vertebrates have demonstrated the integral roles of immune cells in DENV and ZIKV infection, dissemination, and pathogenesis^32–38^, similar studies of mosquito immune cells in arbovirus infection have been constrained by the lack of genetic tools available to manipulate these hemocyte populations. Therefore, previous studies in mosquitoes have been largely observational^23–25,27^ or have relied on methods to impair phagocytic function^26^. To overcome many of these limitations, we have pioneered the use of clodronate liposomes to deplete phagocytic immune cells across arthropod systems^39–41^. Herein, we similarly utilize clodronate liposomes to deplete phagocytic hemocytes in *Ae. aegypti* to determine their respective role in DENV and ZIKV infection. Through our experiments, we demonstrate that phagocytic granulocytes are the hemocyte subtype predominantly infected by arboviruses and that their depletion attenuates virus dissemination. Additional transfer experiments of either acellular hemolymph or phagocytic granulocytes suggest that granulocytes are the primary hemolymph component required to transfer a virus infection. Together, these findings demonstrate the importance of hemocytes as a tropism for virus infection that enhances virus dissemination in the mosquito host.

## Results

### Granulocyte depletion does not influence midgut DENV or ZIKV titers

Mosquito hemocytes have integral roles in innate immunity, yet their roles in arbovirus infection have not been adequately addressed. To approach this question and to determine the influence of granulocytes on midgut virus infection, we employed recently established methods of chemical ablation using clodronate liposomes to selectively deplete phagocytic granulocytes^39–42^. As previously described^40^, *Ae. aegypti* mosquitoes were injected with clodronate liposomes (CLD) to deplete granulocyte populations or with control (LP) liposomes. The effects of CLD treatment were long lasting, with a significant reduction in the percentage of granulocytes when evaluated for more than 10 days post-blood feeding (11 days post-injection) (**Fig. S1a**). Of note, the percentage of granulocytes decreased over time in the LP-treated mosquitoes (**Fig. S1b**), potentially due to an age-related decline in immune function^43^. Despite this, CLD treatment resulted in a steady decrease in the percentage of granulocytes over the entire time examined, such that granulocytes never recovered following CLD treatment (**Fig. S1a**). This suggests that phagocyte depletion is permanent for the life of the mosquito.

To determine the impact of phagocyte depletion on virus infection, mosquitoes were treated with LP or CLD, then challenged with DENV or ZIKV via a blood meal delivered using an artificial membrane system (**Fig. 1a**). Infection outcomes were assessed in dissected midgut samples at 7 days post-infection to determine DENV (**Fig. 1b**) or ZIKV (**Fig. 1c**) viral titers by qRT-PCR. Following CLD treatment, no differences in DENV midgut titers or midgut infection prevalence were observed between LP-or CLD-injected groups (**Fig. 1b**). In addition, LP or CLD treatment similarly had no effect on midgut ZIKV titers, although our data display a significant decrease in infection prevalence in CLD-treated mosquitoes (**Fig. 1c**). These results suggest that phagocytic granulocyte depletion does not have a significant impact on the intensity of midgut flavivirus infection, although significant differences in ZIKV infection prevalence indicate that granulocytes may affect other yet undescribed aspects of midgut physiology that influence the susceptibility to infection in specific virus-vector combinations.

**Figure 1.**
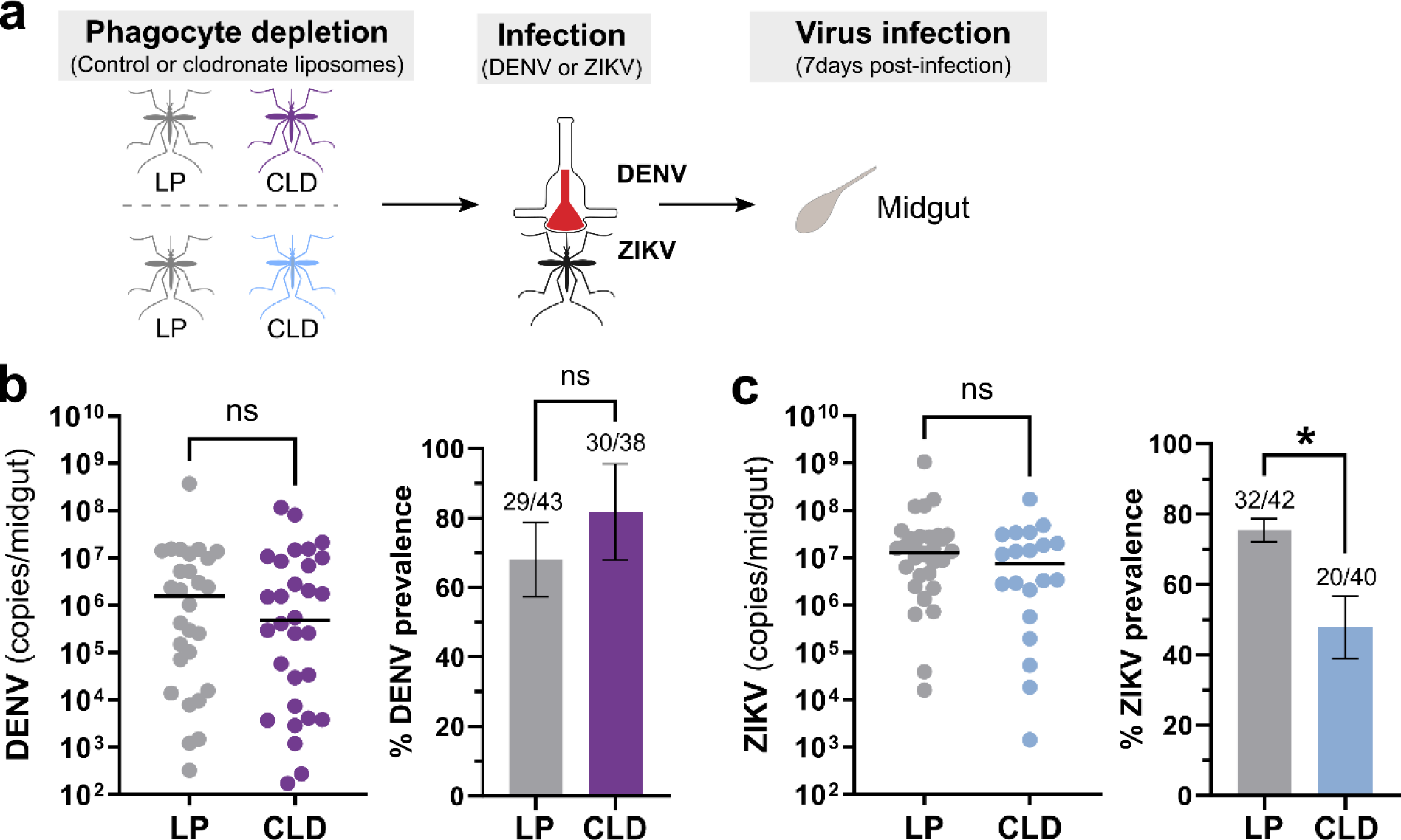
Effects of phagocyte depletion on DENV and ZIKV midgut infection. Overview of experiments performed in *Ae. aegypti* where adult female mosquitoes were injected with either control (LP) or clodronate liposomes (CLD) to examine the effects of phagocytic granulocyte depletion on midgut virus infection (**a**). Approximately 24 hours post-injection, mosquitoes were orally infected with DENV or ZIKV, then midguts were dissected at 7 days post-infection to determine infection outcomes. Viral midgut titers and infection prevalence were determined for DENV (**b**) or ZIKV (**c**) by qRT-PCR, with each dot representing the viral titer of each individual midgut, with the median marked by the black line. The infection prevalence (number of infected mosquitoes of the total analyzed) is displayed in bar graphs as the mean ±SEM. All infection data were pooled from three independent experiments. Viral titers were analyzed using a two-tailed Mann-Whitney test, while prevalence data were examined using a two-sided Fisher’s exact test. Asterisks denote significance (*, *P* < 0.05); ns, not significant.

### Virus primarily infects phagocytic granulocyte populations of mosquito hemocytes

Previous studies demonstrate that arboviruses are able to infect mosquito hemocytes^23–27^, with both prohemocytes^27^ and granulocytes^25,26^ implicated as the predominant hemocyte subtype infected by virus. To further investigate this immune cell specificity for virus infection, DENV-infected mosquitoes (10 days post-infection) were injected with fluorescent beads to enable the identification of phagocytic granulocyte populations as previously described^39,41,42,44^. Following perfusion, hemocytes were examined by IFA for the presence/absence of DENV and the uptake of the fluorescent beads. Similar to previous studies^23,26,27^, DENV readily infected mosquito hemocytes (**Fig. 2a**). While approximately 80% of observed hemocytes were positive for DENV, ∼70% of hemocytes were positive for both DENV and fluorescent beads which are indicative of phagocytic granulocyte populations (**Fig. 2b**). Without other reliable cell markers for *Ae. aegypti* hemocytes it is unclear if the remaining DENV positive cells that did not contain fluorescent beads (∼6% of cells) are prohemocytes, oenocytoids, or granulocytes that failed to uptake a fluorescent bead. These results provide conclusive evidence that phagocytic granulocytes comprise the majority of virus-infected hemocytes and indicate that phagocytic granulocyte populations may be important for virus dissemination.

**Figure 2.**
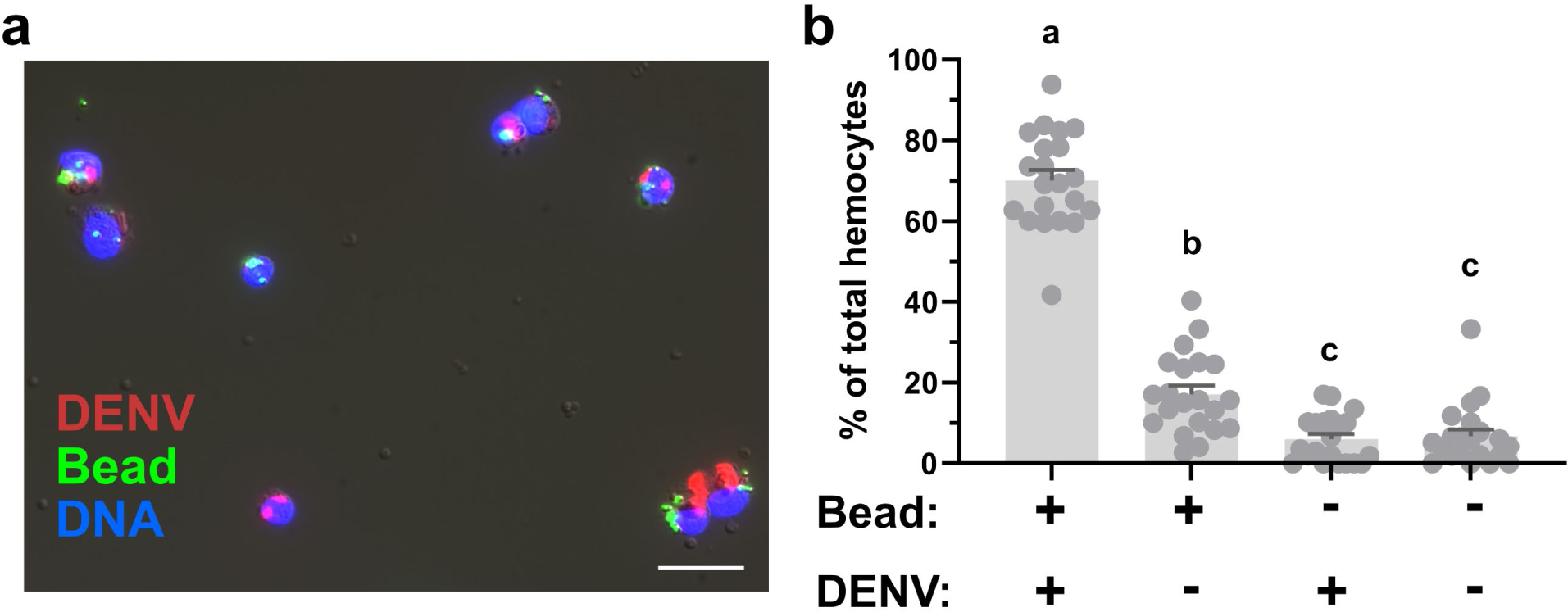
Immunolocalization of DENV in phagocytic granulocytes. Hemocytes were perfused from DENV-infected mosquitoes 10 days post-infecton after injection with fluorescent beads (green). Following fixation, virus localization was examined by immunofluorescence using an anti-DENV monoclonal antibody (clone 3H5-1) followed by an Alexa Fluor 568 goat anti-mouse IgG secondary antibody (red; scale bar: 10 μm) and mounted in ProLong®Diamond Antifade mountant with DAPI (blue) (**a**). To examine the abundance of virus in phagocytic granulocyte populations, the presence/absence of DENV was quantified in the context of the presence/absence of beads used to identify phagocytic granulocyte populations. (**b**) For each experimental outcome, the percentage of the total number of hemocytes for each phenotype are displayed per individual mosquito (dots, n=21) and displayed as the mean ±SEM. Data were pooled from two independent biological experiments. Statistical analysis was performed using Kruskal-Wallis with a Dunn’s multiple comparison test in GraphPad Prism. Letters denote statistically significant differences between samples.

### Phagocytic granulocytes adhere to tissues involved in arbovirus transmission

Given the large proportion of circulating hemocytes infected by virus (**Fig. 2**) and the propensity for hemocytes to adhere to multiple mosquito tissues^22^, the adherence of virus-infected hemocytes to other mosquito tissues may promote virus dissemination. Arguably, the two most important tissues for mosquito arbovirus transmission are the salivary glands, which must become infected for transmission via saliva to a vertebrate host during a blood meal, and the ovaries, which are essential for vertical transmission of virus to offspring. However, the ability of hemocytes to attach to either the salivary glands or ovaries has not been previously characterized. For this reason, we injected mosquitoes with CM-DiI to selectively label hemocytes^44,45^ and fluorescent beads to further identify phagocytic granulocytes as in earlier experiments (**Fig. 2**). We then examined dissected ovaries and salivary glands from mosquitoes 7 days post-infection for the presence of attached phagocytic granulocytes to these tissues using fluorescent microscopy. In both DENV-(**Fig. 3a**) and ZIKV-infected mosquitoes (**Fig. 3b**), we observed phagocytic hemocytes attached to the ovaries and salivary glands, suggesting that adherent granulocytes may facilitate the transport of virus to these secondary tissues required for transmission.

**Figure 3.**
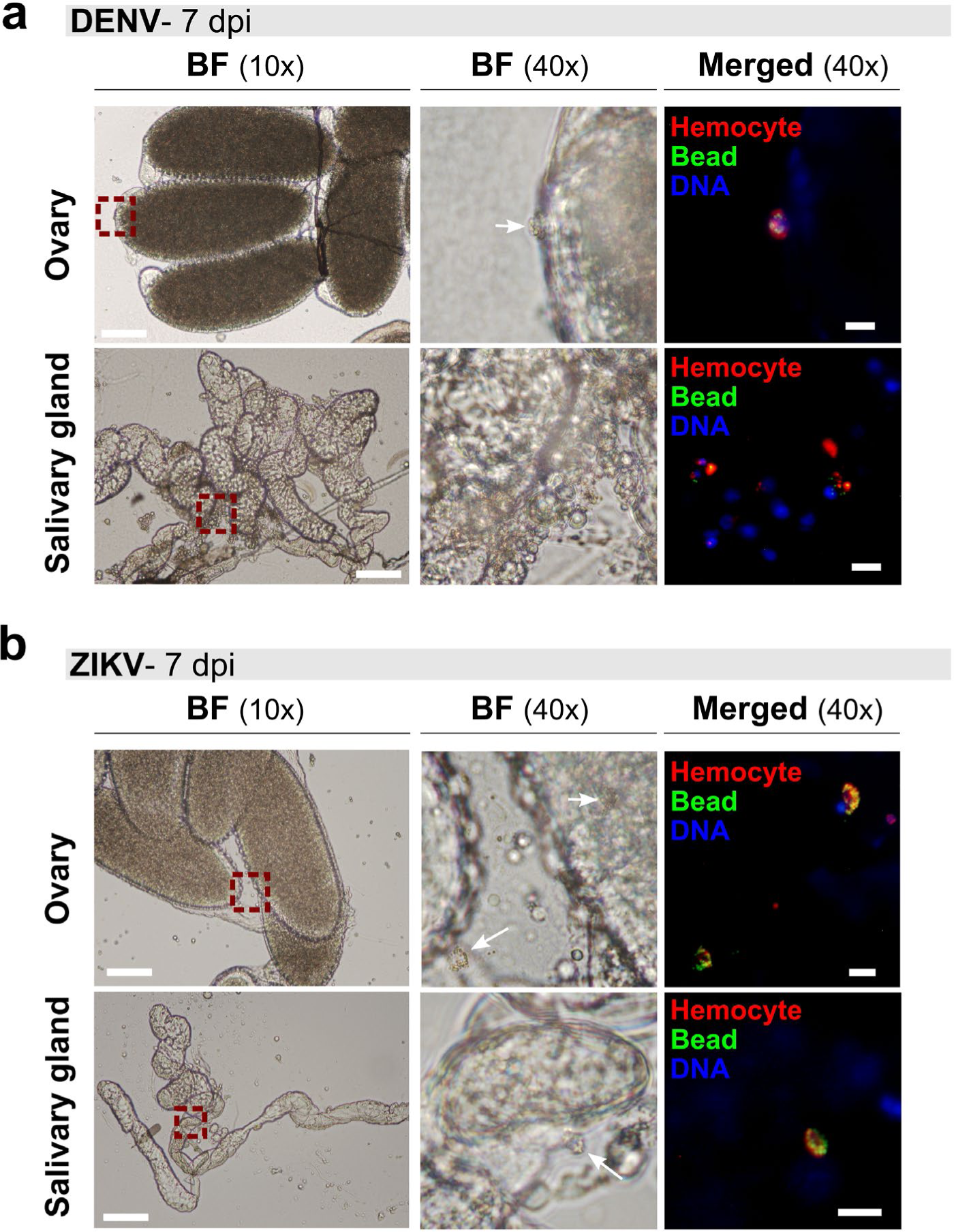
Phagocytic granulocytes associate with the salivary glands and ovaries of virus-infected mosquitoes. To examine hemocyte attachment to mosquito tissues, DENV-(**a**) or ZIKV-infected (**b**) mosquitoes were injected with CM-DiI (red) and fluorescent beads (green) at 7 days post-infection to identify phagocytic granulocyte populations. Following staining *in vivo*, ovary and salivary gland tissues were dissected to examine granulocyte attachment to each respective tissue and mounted using ProLong®Diamond Antifade mountant with DAPI (blue). Red dashed line boxes denote the field of view at 40x magnification, with white arrows used to indicate attached phagocytic hemocytes in the bright field (BF) image where applicable. Scale bars denote 100 μm for 10x images and 10 μm for 40x images.

### Granulocyte depletion attenuates virus dissemination

Since granulocytes become infected with virus (**Fig. 2**) and can attach to the ovaries and salivary glands (**Fig. 3**), we performed experiments to examine the role of hemocytes in virus dissemination. To bypass any potential influence of phagocyte depletion on midgut infection outcomes, mosquitoes were first challenged with DENV or ZIKV, then treated with control or clodronate liposomes three days post-infection (**Fig. 4a**). The influence of phagocyte depletion on virus dissemination was examined in peripheral tissues by measuring virus infection in legs, ovaries, and salivary glands (**Fig. 4a**). DENV RNA titers were significantly reduced in the legs of CLD-treated mosquitoes at 8, 10, and 12 days post-infection (**Fig. 4b**). In addition, there was a significant decrease in the prevalence of DENV infection following granulocyte depletion at day 8 post-infection in the legs, days 8 and 10 post-infection in the ovaries, and day 10 post-infection in the salivary glands (**Fig. 4c**). Similarly, for ZIKV-infected mosquitoes, CLD treatment significantly reduced ZIKV titers in the legs on days 8 and 10 post-infection (**Fig. 4d**), and significantly decreased the prevalence of infection on both day 8 and 10 post-infection in the legs as well as day 8 post-infection in the ovaries and salivary glands (**Fig. 4e**). Together, these results indicate that phagocyte depletion delays virus dissemination from the midgut to the legs, ovaries, and salivary glands, providing strong evidence that granulocytes promote DENV and ZIKV dissemination to peripheral mosquito tissues.

**Figure 4.**
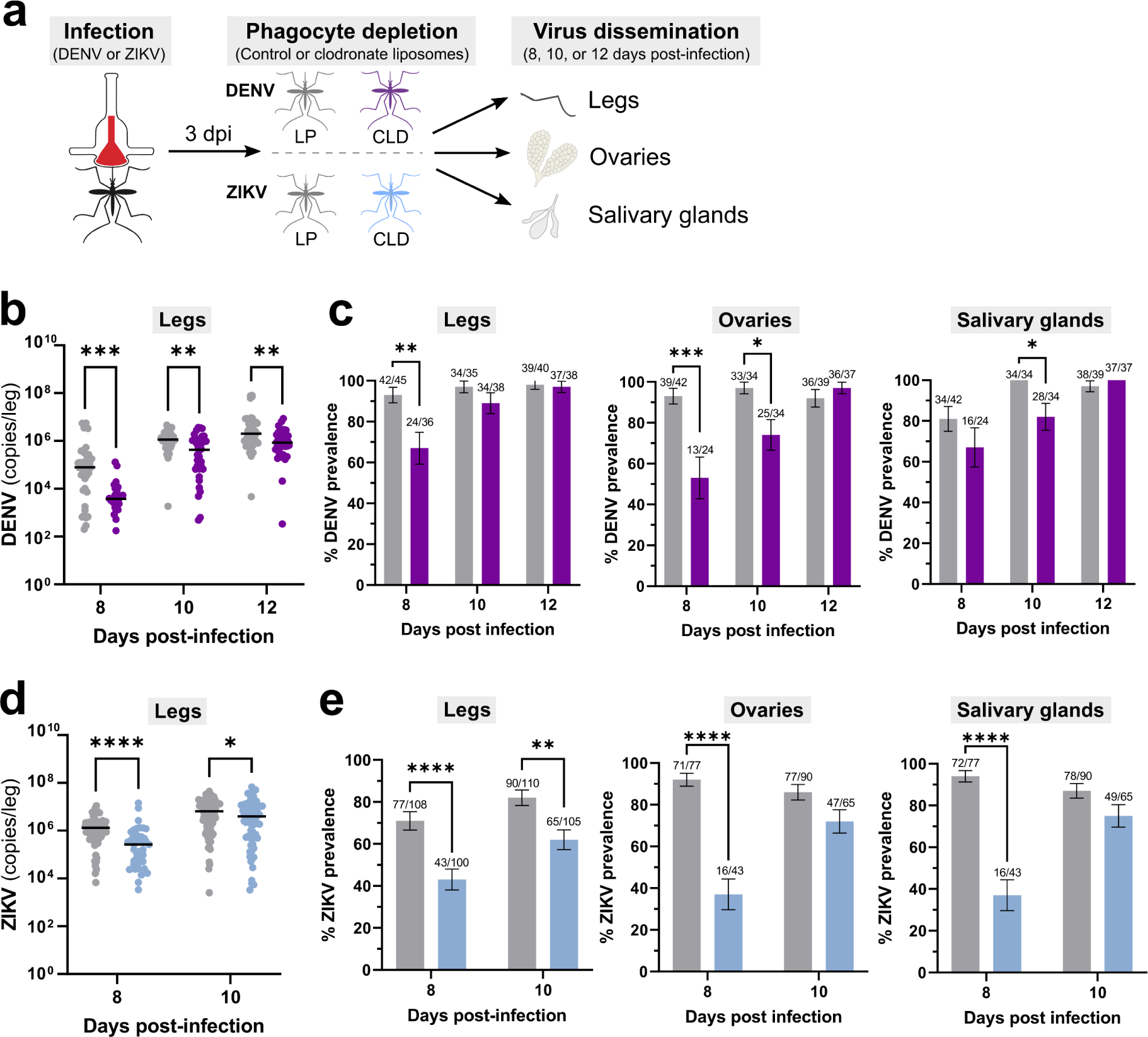
Phagocytic granulocyte depletion impairs virus dissemination to the mosquito legs, ovaries, and salivary glands. (**a**) Overview of virus dissemination experiments. After oral challenge with DENV or ZIKV, mosquitoes were injected with control (LP) or clodronate liposomes (CLD) at 3 days post-infection. Virus dissemination was examined in legs, ovaries, and salivary glands at 8, 10, and 12 days post-infection for DENV (**b** and **c**), or 8- and 10-days post-infection for ZIKV (**d** and **e**). Viral RNA titers are shown for the legs at each respective time point (**b** and **d**), with each dot representing the viral titer of each individual sample and the median marked by the black line. The infection prevalence (number of infected tissues of the total analyzed) is displayed for the legs, ovaries, and salivary glands in bar graphs as the mean ±SEM (**c** and **e**). Infection data were pooled from three independent experiments for DENV and two independent experiments for ZIKV. Viral titers were analyzed using a Mann-Whitney test, with prevalence data analyzed using a Fisher’s exact test. Asterisks denote significance (*, *P* < 0.05; **, *P*<0.01; ***, *P* <0.001, ****, *P* <0.0001); ns, not significant.

### Phagocytic granulocytes transmit virus to uninfected tissues

While virus clearly infects mosquito granulocytes (**Fig. 2**) and the depletion of these immune cells attenuates virus dissemination (**Fig. 4**), it remains unclear whether virus-infected granulocytes represent a viable tropism for virus and if they are able to directly confer an infection to other mosquito tissues. Moreover, the relative contributions of virus-infected hemocytes in virus dissemination compared to free virus circulating in the mosquito hemolymph has not been previously evaluated. To approach these questions, we performed experiments transferring either acellular or cellular fractions of perfused hemolymph from virus-infected mosquitoes to naïve mosquitoes (**Fig. 5a**). Hemolymph was perfused from DENV- or ZIKV-infected mosquitoes at 10 days post-infection, then separated by centrifugation into supernatant (SUP) and cellular fractions (CELL) prior to injection into naïve mosquitoes^46,47^ (**Fig. 5a**). When whole mosquitoes were evaluated at 1, 2, or 4 days post-transfer for DENV or 4 days post-transfer for ZIKV, mosquitoes receiving the cellular fraction had a significantly higher prevalence of DENV (**Fig. 5b**) or ZIKV infection (**Fig. 5c**) when compared to mosquitoes receiving the acellular hemolymph supernatant fraction. Of note, very few mosquitoes injected with the acellular hemolymph supernatant (∼5%) developed an infection (**Fig. 5b** and **5c**), although there were no significant differences in viral genome copy numbers between acellular hemolymph and cell fractions in DENV or ZIKV experiments (**Fig. S2**). To determine if these observations are due to differences in the infectiousness of virus isolates obtained from the supernatant or cell fractions, both isolates were added to cultures of C6/36 cells with infection outcomes monitored at 2- and 3-days post-infection (**Fig. S3**). While DENV cell fractions developed a productive infection, virus collected from the supernatant fraction resulted in a significantly attenuated infection (**Fig. S3a**). In contrast, both the supernatant and cell fractions for ZIKV displayed similar infection outcomes (**Fig. S3b**) despite the cell fraction resulting in a higher prevalence of infection in mosquitoes during transfer experiments (**Fig. 5c**). While we observe some differences between *in vitro* and *in vivo* infectivity of the ZIKV acellular and cellular fractions, these data strongly suggest that hemocytes are the primary component of mosquito hemolymph that is able to promote virus dissemination.

**Figure 5.**
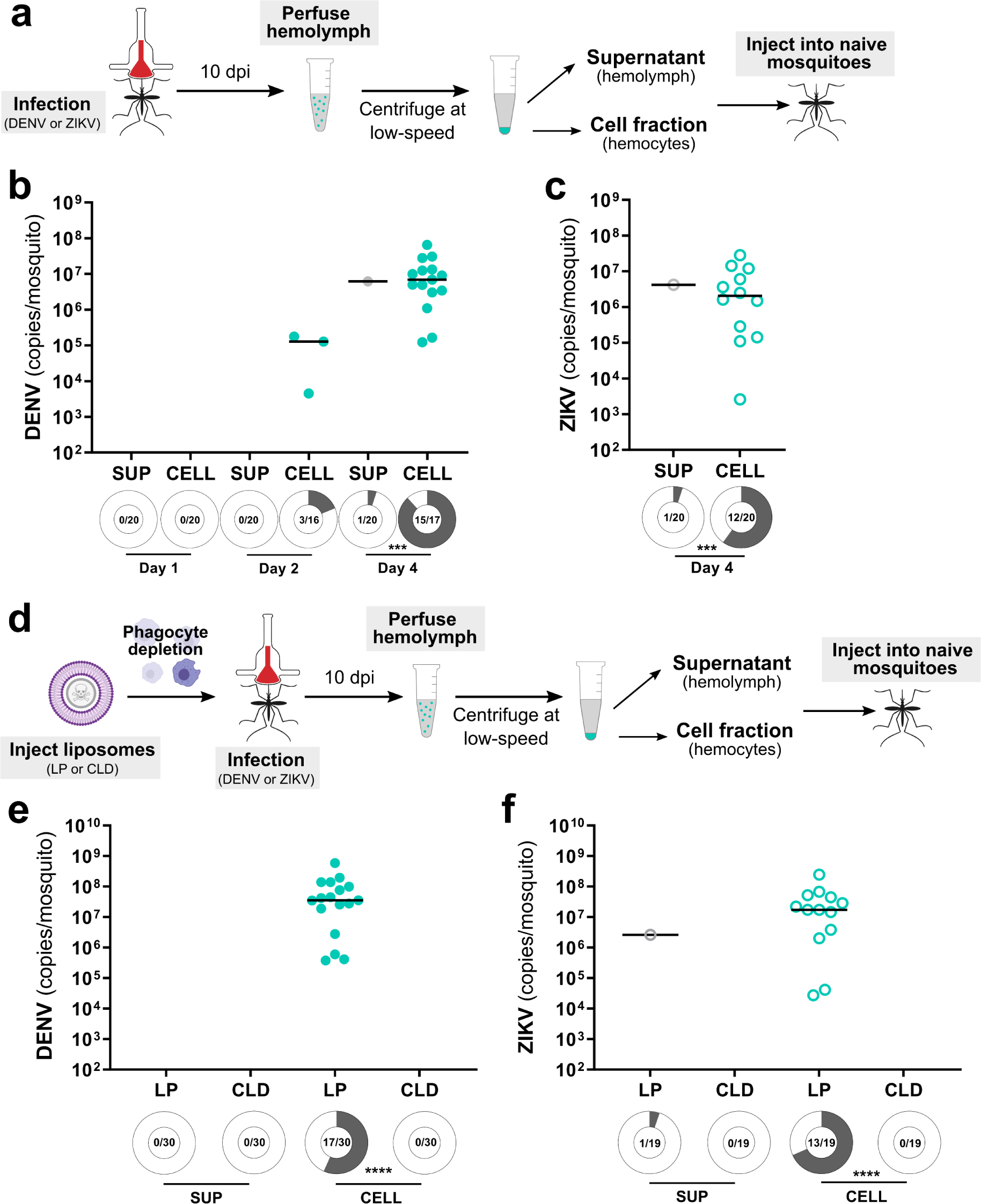
Cellular fractions of the hemolymph more efficiently transfer a virus infection to naïve mosquitoes. To assess the respective infectivity of the cellular and acellular fractions of mosquito hemolymph, *Ae. aegypti* were first infected with DENV or ZIKV, then the hemolymph was perfused from mosquitoes at 10 days post-infection (**a**). Hemolymph was separated into hemocyte-containing cell (CELL) and acellular hemolymph supernatant (SUP) fractions using low-speed centrifugation and then injected into naïve mosquitoes (**a**). Whole mosquitoes were evaluated for viral titer and infection prevalence by qRT-PCR at 1, 2, and 4 days post-injection for DENV (**b**) and 4 days post-injection for ZIKV (**c**). Each dot represents the viral titer of individual samples with the median displayed by the black line. Similar experiments were performed where phagocytic granulocytes were depleted with clodronate liposomes (CLD) prior to DENV or ZIKV infection to confirm that granulocytes are required for the transfer of virus infection in the cellular fractions of the mosquito hemolymph (**d**). Empty liposomes (LP) serve as a control. Whole mosquitoes were assessed for viral titer and infection prevalence at 4 days post-injection for both DENV (**e**) and ZIKV (**f**). Due to the low numbers of virus positive samples in certain control and experimental conditions, viral titers were not directly compared. Infection prevalence data were analyzed using a Fisher’s exact test. Shaded areas of circles under each experimental condition correspond to infection prevalence, with the number infected of the total number of mosquitoes examined displayed for each treatment. Data were pooled from two independent experiments. Asterisks denote significance (***, *P* <0.001; ****, *P* < 0.0001).

With our previous results suggesting that phagocytic granulocytes are an important factor in virus dissemination (**Figs. 2-5**), we repeated our transfer experiments in the context of phagocyte depletion to determine if granulocytes are the cell type responsible for the successful viral infection of the naïve mosquito host following transfer (**Fig. 5d**). Similar to our earlier transfer experiments (**Fig. 5b** and **5c**), transfer of the cell fraction from control mosquitoes injected with LP produced a virus infection in naïve mosquitoes when examined 4 days post-transfer for either DENV (**Fig. 5e**) or ZIKV (**Fig. 5f**). In contrast, when samples originated from CLD-treated mosquitoes, the transfer of virus was completely abolished for both DENV (**Fig. 5e**) and ZIKV (**Fig. 5f**). Similar to previous results (**Figs. 5b** and **5c**), the supernatant fraction was unable to efficiently transfer a viral infection to naïve mosquitoes (**Figs. 5e** and **5f**). Taken together, these results provide strong evidence that granulocytes are capable of acquiring and disseminating flaviviruses to mosquito tissues required for arbovirus transmission.

## Discussion

While immune cells are instrumental to DENV and ZIKV infection and dissemination in vertebrate systems^32–38^, we still lack a critical understanding of the mechanisms that drive virus infection and dissemination in the mosquito host. These efforts have previously been hindered by the lack of genetic tools to manipulate mosquito hemocyte populations, thereby limiting our ability to study these immune cells and their contributions to arbovirus infection. By taking advantage of recently described methods to deplete phagocytic hemocyte populations in arthropods^39–41^, we demonstrate that phagocytic hemocytes support arbovirus infection and facilitate virus dissemination to important tissues such as the salivary glands and ovaries in the mosquito *Ae. aegypti*.

Our phagocyte depletion experiments suggest that DENV and ZIKV titers in the mosquito midgut are not directly influenced by the presence or absence of phagocytic granulocyte populations. However, the reduced prevalence of midgut ZIKV infection following granulocyte depletion suggests that these immune cells may still contribute to the success of midgut infection, potentially through the regulation of epithelial homeostasis^48,49^. Similar pro-viral effects of phagocytic granulocytes on midgut virus infection have been previously described, yet in contrast to our own study, previous studies show reduced DENV and ZIKV midgut titers when populations of phagocytic granulocytes are functionally overloaded via bead injection^26^. While both methodologies target granulocytes, bead injection may also promote other unintended cellular or humoral responses that reduce midgut infection or virus replication, as evidenced by an increase in midgut-associated hemocytes after bead injection^26^. While further study is required to fully delineate the complex role of hemocytes in midgut viral infection and replication, our results suggest that the overall influence of granulocytes on midgut infection are minimal and perhaps virus- or titer-dependent.

Similar to previous studies^12,23–26^, we observed virus localization in circulating hemocytes collected from perfused hemolymph. However, there have been disagreements as to the immune cell subtypes infected by virus, with previous studies implicating either prohemocytes^27^ or granulocytes^25,26^. Through immunofluorescence experiments pairing virus infection with the uptake of fluorescent beads, we demonstrate that the majority of virus-infected hemocytes are phagocytic granulocytes, indicating that these immune cell populations are a focal point for virus infection in the hemolymph. With previous experiments supporting the existence of multiple granulocyte subtypes^39,42,50,51^, additional experiments are required to determine if specific subpopulations of granulocytes differ in their susceptibility to virus infection.

While our work and that of others has demonstrated that virus can infect the hemocytes of mosquitoes and other insects^12,23–26^, the exact mechanism and location of hemocyte virus acquisition remains unknown. Previous work has shown that hemocytes adhere to multiple mosquito tissues^45^ and are recruited to the midgut following virus infection^26^. Therefore, it is possible that hemocytes acquire an infection after direct contact with the midgut basal lamina, where mature virions are concentrated^15^. Additional studies have demonstrated that arboviruses infect the tracheal network associated with the midgut epithelium^23,52,53^, potentially providing other routes of infection similar to that required for hemocyte infection and baculovirus dissemination in Lepidoptera^12,54^. Alternatively, hemocytes may also acquire infection through the uptake of free virus in the hemolymph. Given these multiple possible routes of infection, further investigation is required to determine the mechanisms by which virus infects mosquito hemocytes.

When clodronate depletion experiments were performed after virus midgut infection to bypass any effect on midgut infection outcomes, we found that phagocytic granulocyte populations have an integral role in the dissemination of virus. Phagocyte depletion reduced infection prevalence and viral titers in the legs, a tissue routinely used as an indicator of transmission potential^55^. In addition, phagocyte depletion lowered the prevalence of DENV and ZIKV in the ovaries and salivary glands, the infection of which can respectively lead to vertical transmission to offspring or transmission to a vertebrate host during blood-feeding. These data are further supported by our immunofluorescence experiments confirming that phagocytic granulocytes attach to both ovaries and salivary glands in DENV- and ZIKV-infected mosquitoes. Together, these data suggest that phagocytic granulocytes make significant contributions to DENV and ZIKV dissemination in the mosquito and likely enhance virus transmission to both the vertebrate host and invertebrate offspring. Despite this seemingly pro-viral role for phagocytic granulocytes, it should be noted that the loss of these immune cell populations only attenuates virus dissemination. At present, it is unclear if this is due to incomplete phagocyte depletion using clodronate liposomes, the presence of free virus in the hemolymph, or if additional, yet unexplored, mechanisms of virus dissemination occur in the mosquito host.

Through the use of transfer experiments using either acellular or cellular hemolymph fractions, we demonstrate that virus-infected hemocytes are capable of conferring a productive viral infection to a naïve host. Through clodronate depletion, we specifically implicate phagocytic granulocytes in this virus transfer, providing strong support for the importance of these immune cells in virus dissemination. In contrast, we found that the acellular hemolymph was far less efficient in transferring a productive infection to naïve mosquitoes despite containing comparable virus loads when measured by qRT-PCR. This suggests that additional unknown factors restrict the infectivity of free virus in the hemolymph and argue that hemocyte virus infection enhances the success of virus dissemination. Therefore, we believe that virus-infected granulocytes behave as a “trojan horse”, protecting intracellular virus from antiviral factors present in the hemolymph, while serving as a vehicle to deliver virus to target tissues, such as the salivary glands and ovaries.

Similar roles in arbovirus infection have also been described for vertebrate immune cell populations. While tissue-resident dendritic cells (such as Langerhans cells) provide sites of initial arbovirus replication and local dissemination^56–59^, monocytes and monocyte-derived macrophages transport virus through the circulatory system to further enhance virus replication, persistence, and dissemination^38,57,60^. This includes comparable function of immune cells as “trojan horses” in vertebrate systems, where monocyte/macrophage populations mediate viral entry and dissemination in the central nervous system^60^. With monocytes/macrophages performing as professional phagocytes similar to that of mosquito granulocytes, we believe that this represents an exciting parallel between vertebrate and invertebrate systems in the manner by which virus exploit immune cell populations for their own replication and propagation.

In addition to the roles of mosquito hemocytes in virus dissemination outlined in this study, hemocytes also contribute to anti-viral immunity. Previous studies in *Ae. aegypti* demonstrate that impairing hemocyte function by overloading these cells with beads increased viral titers in whole mosquitoes injected with ZIKV or DENV^26^. This implies that phagocytic activity in hemocytes restricts virus dissemination. Similar observations have also been described for *Drosophila*, where the uptake of virus RNA by hemocytes mediates a systemic RNAi anti-viral response that limits virus replication^31^. Therefore, it is likely that hemocytes play a complex role in viral infection and may be involved in both anti-viral and pro-viral activities that may vary among specialized hemocyte sub-populations.

While our study examines the influence of hemocytes in flavivirus (DENV and ZIKV) dissemination, the role of hemocytes in virus infection and dissemination may differ for other arboviruses. For instance, alphaviruses and flaviviruses have different strategies for replication and assembly^61–63^ where alphaviruses typically replicate faster in the mosquito host. For example, chikungunya virus is able to quickly replicate and escape the midgut as early as 24 h after an infectious blood meal^15,64^, contrasting the approximate three days required for the earliest escape of ZIKV from the midgut^16^. As a result, alphaviruses may be able to exploit transient damage to the basal lamina resulting from blood-feeding prior to its repair, potentially enabling free virus to escape into the hemolymph and disseminate to other mosquito tissues without the need of hemocytes. However, another alphavirus, Sindbis virus, has been shown to infect and replicate within hemocytes^25^, implying that hemocytes may have similar roles in alphavirus dissemination. Further investigation is required to more precisely determine whether the roles of hemocytes in arbovirus infection are conserved for all mosquito-borne arboviruses.

In summary, the results of this study support three main conclusions. First, phagocytic granulocytes are focal points for virus infection and attach to mosquito tissues involved in arbovirus transmission. Second, the depletion of granulocytes via clodronate liposomes delays viral dissemination, implying that granulocytes significantly contribute to the dissemination of virus to tissues such as the salivary glands and ovaries. Third, virus-infected granulocytes are the primary component of the hemolymph able to transfer a viral infection to a naïve mosquito. Taken together, the results of this study converge upon a model for virus dissemination whereby hemocytes acquire virus and subsequently transport an infection via the hemolymph to other mosquito tissues (**Fig. 6**). While the exact mechanisms through which hemocytes acquire and transmit a viral infection remain to be explored, the hemocyte tropism for viruses described here suggest that hemocytes have profound impacts on mosquito vector competence and the extrinsic incubation period for both DENV and ZIKV. The results of this study significantly advance our understanding of virus infection dynamics in mosquitoes that highlight conserved roles of immune cells in virus dissemination.

**Figure 6.**
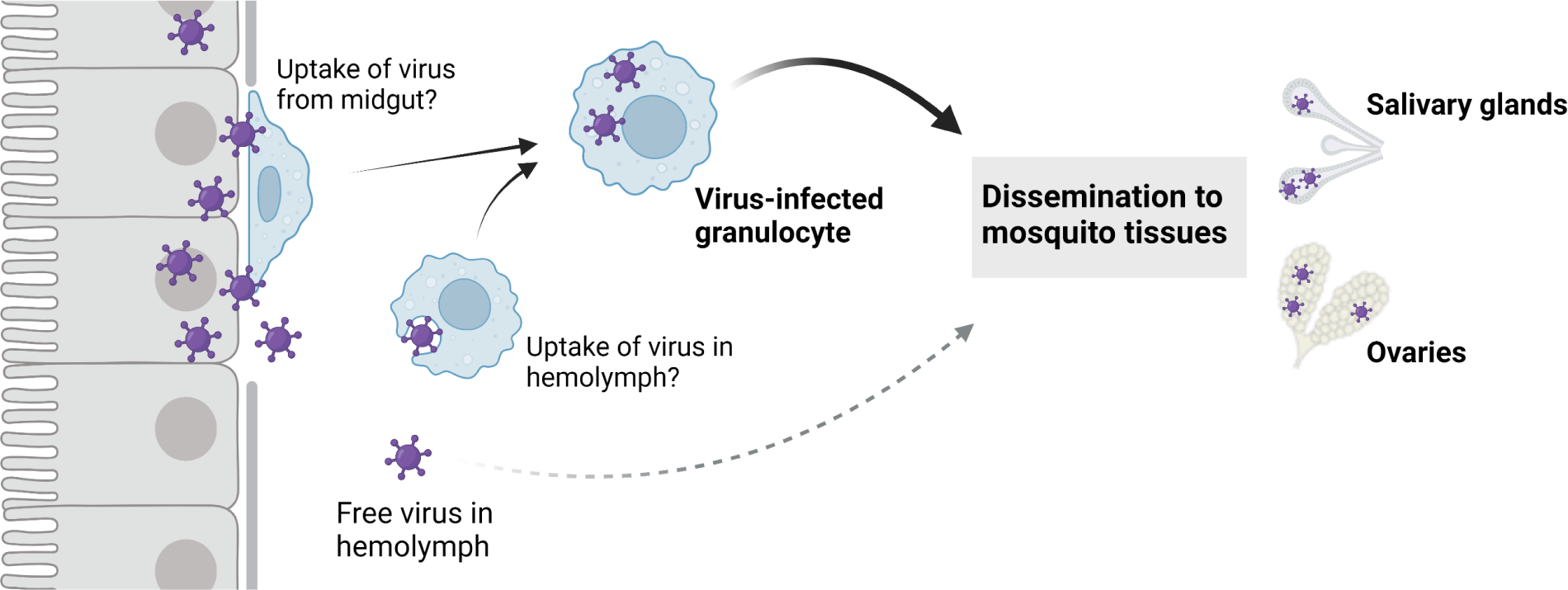
“Trojan horse” model of virus dissemination in the mosquito host. Following virus infection of the mosquito midgut, a subset of phagocytic granulocytes acquire virus either through attachment to the virus-infected midgut, the uptake of free virus present in the mosquito hemolymph, or the attachment to other infected mosquito tissues such as the trachea (not shown). Through the movement of these immune cells in the hemolymph and their ability to adhere to mosquito tissues, virus-infected granulocytes display increased efficiency to disseminate virus to the salivary glands and ovaries when compared to free virus present in the hemolymph. This suggests that these virus-infected granulocytes act as a “trojan-horse” to enhance virus dissemination.

## Methods

### Cell culture and virus isolates

*Ae. albopictus* C6/36 cell maintenance and virus culture were performed at Iowa State University (ISU) and the Connecticut Agricultural Experiment Station (CAES) with slight modifications. At ISU, *Ae. albopictus* C6/36 cells were maintained in 25 cm^2^ flasks at 28°C in L-15 media supplemented with 10% fetal bovine serum (FBS), 4 mM L-glutamine, 100 units/ml penicillin, and 100 µg/ml streptomycin. At CAES, *Ae. albopictus* C6/36 cells were grown in 75 cm^2^ flasks using minimum essential medium containing 2% FBS, 1X non-essential amino acids, 100 U/ml penicillin, and 100 μg/ml streptomycin. Virus infection experiments at both locations were performed with DENV serotype 2 (125270/VENE93; GenBank Accession No. U91870) and ZIKV (PRVABC59; GenBank Accession No. KU501215). To infect cells with virus, growth media was removed and replaced with 2 ml L-15 media supplemented with 2% fetal bovine serum, 4 mM L-glutamine, and 250 µl frozen DENV or ZIKV stock, and left at room temperature rocking periodically for 15 minutes. The flask was then placed at 28°C for 45 minutes rocking periodically, after which 3 ml of additional media supplemented with 2% FBS and 4 mM L-glutamine was added. The infection was allowed to proceed until significant cytopathic effects were observed, approximately 4 days for DENV and 5 days for ZIKV. At CAES, 250 µl DENV viral stocks or 250 µl ZIKV were diluted in 3 ml media and used to infect flasks of C6/36 cells at 70-80% confluency. Flasks were placed on a rocking platform with infection media for 1 hr at room temperature before adding 12 ml regular media. Infected cells were incubated at 28°C for 5 days.

### Mosquito rearing and virus infection

All experiments were performed using *Ae. aegypti* (Orlando strain) with slight modifications between performance site locations. At ISU, mosquitoes were reared at 27°C and 80% humidity with a 16:8 light: dark cycle. Larvae were reared on a diet of ground fish flakes (Tetramin, Tetra), while adult mosquitoes were maintained on a 10% sucrose solution. Mosquito colonies were maintained via artificial membrane feeding using commercial defibrinated sheep blood (HemoStat Laboratories) for egg production. At CAES, mosquitoes were reared at 27°C and 80% humidity with a 14:10 light: dark cycle. Larvae were reared with a 3:2 mix of powdered liver:yeast, while adults were maintained on 10% sucrose. For virus infection experiments at both locations, adult female mosquitoes (3-5 days after adult emergence) were starved overnight and challenged using a 1:1 mixture of defibrinated sheep blood and fresh DENV or ZIKV culture using a Hemotek artificial membrane feeding system. Virus titers for infection experiments are displayed in **Table S1**. After feeding, mosquitoes were immediately cold anesthetized to allow for the selection of blood-fed mosquitoes.

### Midgut infection experiments

To assess the impact of granulocyte depletion on midgut infection outcomes, naïve mosquitoes (3-5 days post-eclosion) were injected with either 69 nl of control liposomes (LP) or clodronate liposomes (CLD) (Encapsula NanoSciences) at a 1:4 dilution as previously^40^. At 24 hours post-injection, mosquitoes were then orally infected with DENV or ZIKV. Midguts were isolated in 1X PBS at 7 days post-DENV or -ZIKV infection, individually homogenized in 400 µl TRIzol Reagent (Invitrogen), and then left at 4°C overnight to maximize RNA yield. RNA isolation was completed using the Direct-zol RNA MiniPrep kit (Zymo Research) following the manufacturer’s protocols. Viral RNA titers were determined by qRT-PCR using the QuantStudio 3 (ThermoFisher) and TaqMan Fast Virus 1-Step Master Mix (ThermoFisher). Reactions were set up according to the manufacturer’s protocols using a 10 µl reaction volume and 25 ng of total RNA template. qRT-PCR reactions were performed using previously described primers for DENV^65^ or ZIKV^66^ (**Table S2**) with the following PCR conditions: 50°C for 5 min, 95°C for 20 sec, and 40 cycles of 95°C for 3 sec, 60°C for 30 sec. For each experiment, a standard curve was calculated using 6 standards ranging from 10^2^ to 10^7^ genome copies per reaction amplified in duplicate on each 96-well plate. Virus genome copy number per reaction was determined by absolute quantification using a standard curve, with the lower limit of detection set at the lowest standard (10^2^ genome copies per reaction).

### Immunofluorescence assays for localization of viruses in hemocytes

Hemocyte immunofluorescence assays (IFAs) were performed as previously described^39,44,67^. Following infection with DENV, mosquitoes were injected with 69 nl of a 2% green fluorescent FluoSpheres solution (in 1X PBS) using a Nanoject III (Drummond Scientific) and incubated at 27°C for 1 h to allow for bead uptake by hemocytes at 10 days post-infection. Hemolymph perfusion was carried out as previously described^39,42,44,67,68^. Briefly, an incision was made in the second to last abdominal segment, and approximately 10 µl of anticoagulant solution (60% [vol/vol] Schneider’s insect medium, 10% FBS, and 30% citrate buffer; 98 mM NaOH, 186 mM NaCl, 1.7 mM EDTA, and 41 mM citric acid, pH 4.5) was used to perfuse hemolymph using a Nanoject III injector onto a multi-well glass slide (MP Biomedicals). After 30 min incubation, cells were fixed with 4% paraformaldehyde for 15 minutes at room temperature (RT) and then washed three times in 1X PBS. Samples were incubated with blocking buffer (0.1% Triton X-100, 1% BSA in 1X PBS) for 24 h at 4°C, then with an anti-DENV monoclonal antibody (clone 3H5-1; BEI Resources) at a 1:200 dilution in blocking buffer overnight at 4°C. After washing three times in 1X PBS, an Alexa Fluor 568 goat anti-mouse IgG (1:500, Thermo Fisher Scientific) secondary antibody in blocking buffer was added for 2 h at RT. Slides were rinsed three times in 1X PBS and mounted with ProLong®Diamond Antifade mountant with DAPI (Life Technologies). Images were examined by fluorescence microscopy (Nikon Eclipse 50i, Nikon). To determine the proportion of phagocytic hemocytes infected by virus, 50 hemocytes from randomly chosen fields were examined for the presence or absence of virus signal and/or beads.

### Hemocyte attachment to mosquito salivary glands and ovaries

At 7 days post-DENV or -ZIKV infection, mosquitoes were injected with 69 nl of a solution containing 100µM Vybrant CM-DiI (ThermoFisher Scientific) and 2% green fluorescent beads in 1X PBS, then incubated under insectary conditions for 30 min. To preserve hemocyte attachment, mosquitoes were injected with 207 nl of 50% ethanol in 1X PBS. Salivary glands and ovaries were then dissected in 1X PBS and mounted using ProLong®Diamond Antifade Mountant with DAPI. The attachment of phagocytic granulocyte populations (cells that are DiI^+^/bead^+^) to salivary glands and ovaries were examined by fluorescence microscopy (Nikon Eclipse 50i, Nikon).

### Virus dissemination experiments

To assess the impact of hemocyte depletion on virus dissemination, mosquitoes were first orally challenged with either DENV or ZIKV, then injected with 69 nl of a 1:4 dilution of LP or CLD as previously described^40^ at 3 days post-infection. To assess DENV dissemination, mosquito legs, ovaries, and salivary glands were harvested at 8-, 10-, and 12-days post-infection. For ZIKV assays, mosquito legs, ovaries, and salivary glands were harvested at 8- and 10-days post-infection. RNA was isolated from dissected tissues with a Mag-Bind Viral DNA/RNA 96 Kit (Omega Bio-tek Inc) and Kingfisher Flex automated nucleic acid extraction device (ThermoFisher Scientific) per the manufacturer’s instructions. Viral RNA titers and infections were assessed via qRT-PCR using an iTaq Universal Probes One-Step Kit (BioRad) in a 20 µl sample volume. Reactions were performed using previously described primers for DENV-2^69^ or ZIKV^66^ (**Table S3**) with the following PCR conditions: 50°C for 30 minutes, 95°C for 10 min and 40 rounds of 50°C for 15 sec and 60°C for 1 min. Viral RNA titers were quantified using a standard curve with a cut-off value of Ct<36. Sample Ct values below this cut-off were considered positive whereas samples with larger Ct values were considered negative. Viral concentrations were corrected for dilution factor. Results were pooled for DENV and ZIKV assays from either three or two independent biological replicates respectively.

### Hemolymph perfusion and hemocyte transfer experiments

To examine the infectivity of the cellular and acellular fractions of the mosquito hemolymph, cellular and cell-free hemolymph fractions were isolated from virus-infected mosquitoes and transferred to naïve mosquitoes similar to previous studies^46^. At 10 days post-infection with DENV or ZIKV, perfused hemolymph was collected using ∼10 µl of transfer buffer (95% Scheider’s medium and 5% citrate buffer) from individual mosquitoes. Pooled samples (n=15) were separated into hemocyte-containing (CELL) and cell-free supernatant (SUP) fractions by centrifugation at 4°C, 2000 g for 5 min. While the SUP fraction was used directly, the CELL fraction was additionally washed in 1X PBS and again centrifuged at 4°C, 2000 g for 5 min. Following centrifugation, the 1X PBS was removed and the resulting CELL pellet was resuspended in an equivalent volume of transfer buffer. Naïve mosquitoes were injected with 207 nl of either the prepared SUP or CELL fractions. Whole mosquitoes were collected at 1, 2, and 4 days post-injection for DENV and 4 days post-injection for ZIKV. Whole mosquitoes were homogenized in 400 µl TRIzol Reagent and RNA isolation was then completed using Direct-zol RNA MiniPrep kit. Viral titers of whole mosquitoes were analyzed by qRT-PCR using the above described protocols for midgut infection experiments using 100 ng of template RNA per reaction.

Additional experiments were performed to examine the effects of phagocyte depletion on the infectivity of cellular and acellular hemolymph fractions. Briefly, naïve mosquitoes were injected with 69 nl of LP or CLD (1:4 dilution) as described above and then challenged with an oral infection of DENV or ZIKV. Hemolymph perfusion, isolation of SUP and CELL fractions, and transfer to naïve mosquitoes was carried out as described above. Viral RNA titers of whole mosquitoes were examined by qRT-PCR 4 days post-transfer.

To confirm the infectivity of the cellular and acellular hemolymph fractions *in vitro*, C6/36 cells were first seeded into a 24-well plate at approximately 1×10^6^ cells per well and allowed to attach for 24 h. Collected SUP and CELL fractions were diluted in 600 µl of L-15 media supplemented with 2% FBS, 4 mM L-glutamine, 100 units/ml penicillin, 100 µg/ml streptomycin and 0.25 µg/ml amphotericin B. Then 300 µl of infectious media from each fraction was added to C6/36 cells in duplicate. To determine the initial titer of each hemolymph fraction, additional SUP and CELL isolates collected from the same group of infected mosquitoes were directly lysed in 400 µl TRIzol Reagent and left at 4°C overnight to maximize yield. Infection outcomes were examined at 2- and 3-days post-infection by resuspending the C6/36 cells by vigorously pipetting up and down, then 100 µl of the cell suspension was added to 900µl TRIzol Reagent. RNA isolation was then completed using Direct-zol RNA MiniPrep kit and viral RNA titers determined by qRT-PCR.

### Hemocyte percentages

To quantify the efficacy of CLD depletion of phagocytic hemocytes, mosquitoes (3-5 days old) were injected with 69 nl of 1:4 diluted LP or CLD as described above. Surviving mosquitoes were challenged with defibrinated sheep blood (Hemostat Laboratory) 24 h post-injection. Hemolymph was perfused at 1, 4, 7, and 10 days post-blood feeding using an anticoagulant solution as previously described^39,44^. The proportion of granulocytes was calculated by analyzing ∼200 hemocytes per individual mosquito based on morphology using a disposable Neubauer hemocytometer slide (C-Chip DHC-N01; INCYTO) as previously described^39,44^.

## Supporting information

Supporting Information containing Figures S1 to S3 and Tables S1 to S3

## Acknowledgements

This work was supported by R21 AI149118 (DEB and RCS) and R01 AI148477 (DEB) from the National Institutes of Health, National Institute of Allergy and Infectious Diseases. This material is based upon work supported by the National Science Foundation Graduate Research Fellowship Program under Grant No. 2336877. Any opinions, findings, and conclusions or recommendations expressed in this material are those of the authors and do not necessarily reflect the views of the National Science Foundation. The monoclonal anti-dengue virus type 2 envelope protein antibody, clone 3H5-1 (NR-2556) was obtained through BEI Resources, NIAID, NIH.

