## Supporting Information containing Figures S1 to S3 and Tables S1 to S3 for "Mosquito immune cells enhance dengue and Zika virus dissemination in *Aedes aegypti*"

### Included supporting information

#### *Supplemental Figures*

**Figure S1.** Stable depletion of granulocytes by clodronate liposome injection.

**Figure S2.** Viral titers of acellular and cellular fractions collected from mosquito hemolymph.

**Figure S3.** *In vitro* infections using acellular and cellular hemolymph fractions.

#### *Supplemental Tables*

**Table S1.** Summary of blood meal virus titers for infection experiments.

**Table S2.** Primers used to assess virus titers in midgut and transfer experiments.

**Table S3.** Primers used to assess virus titers in dissemination experiments.

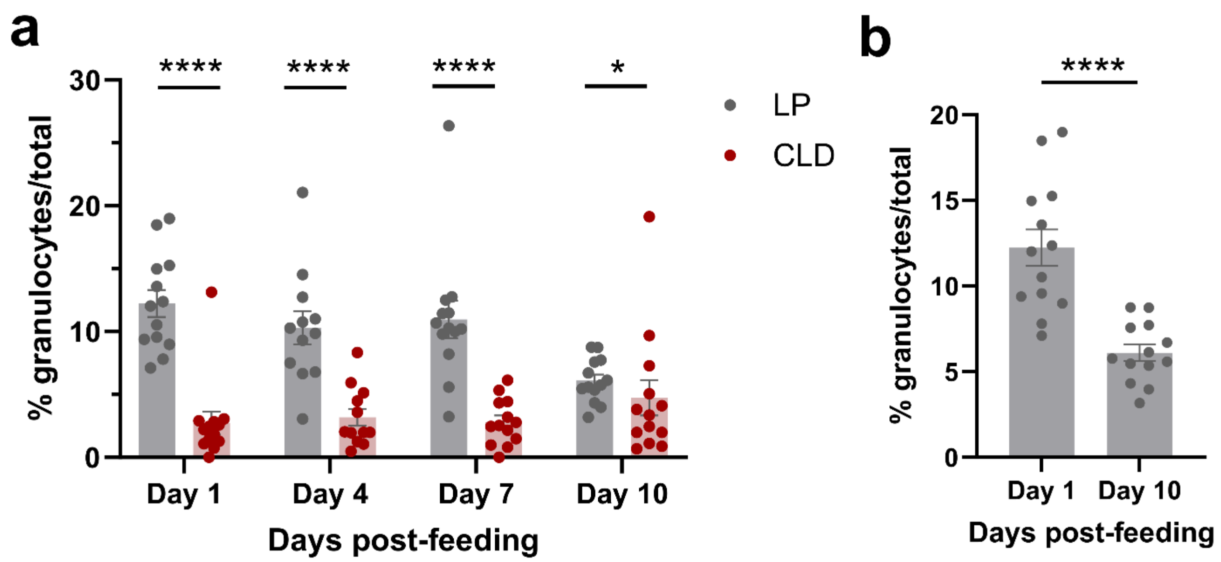

**Figure S1. Stable depletion of granulocytes by clodronate liposome injection.** To validate the effectiveness and duration of granulocyte depletion by clodronate liposomes in blood-fed *Ae. aegypti*, mosquitoes were first injected with either clodronate liposomes (CLD) to deplete phagocytic granulocyte populations or control liposomes (LP) and then blood-fed at 24 hours post-injection. Hemolymph was perfused at days 1, 4, 7, and 10 post-feeding and the percentage of granulocytes of the total hemocyte population were determined using a hemocytometer (**a**). In control LP-injected mosquitoes, the percentage of granulocytes was significantly lower at day 10 when compared to day 1 (**b**), suggesting that there is a decline in the percentage of granulocytes with age. Data were analyzed using a two-tailed Mann-Whitney test. Asterisks denote significance (*, $P$ <0.05; ****, $P$ <0.0001).

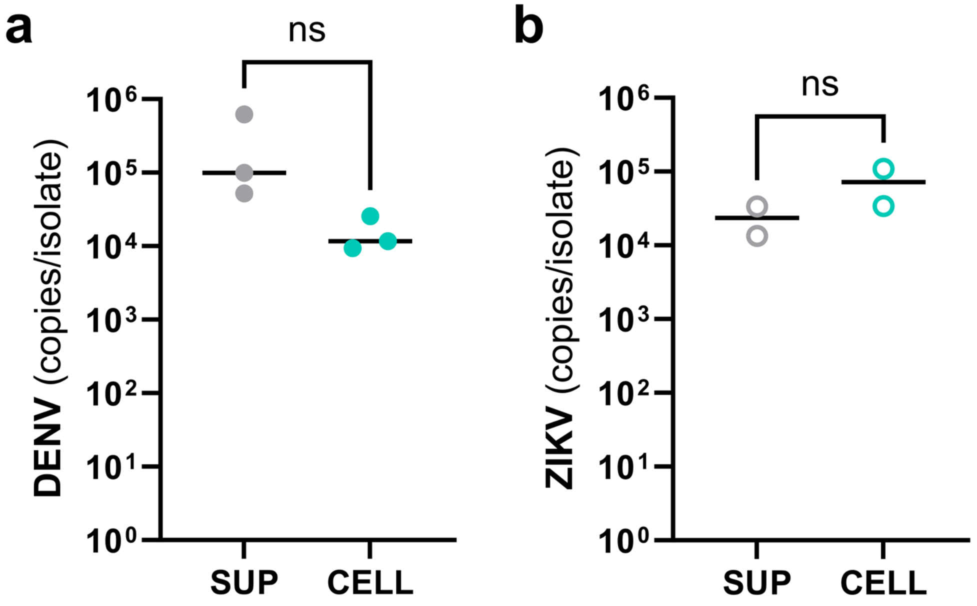

**Figure S2. Viral titers of acellular and cellular fractions collected from mosquito hemolymph.** At 10 days post-infection, hemolymph was perfused from DENV- or ZIKV-infected mosquitoes. Following centrifugation, hemolymph was separated into acellular (SUP) and cellular (CELL) fractions. Total RNA was isolated from each fraction and the viral RNA titer of each fraction taken from DENV- (**a**) or ZIKV-infected mosquitoes (**b**) was determined by qRT-PCR. Each dot represents the viral titer of independent biological replicates with the median displayed by the black line. Data from two (ZIKV) or three (DENV) independent biological experiments were analyzed using a two-tailed Student's t-test. ns, not significant.

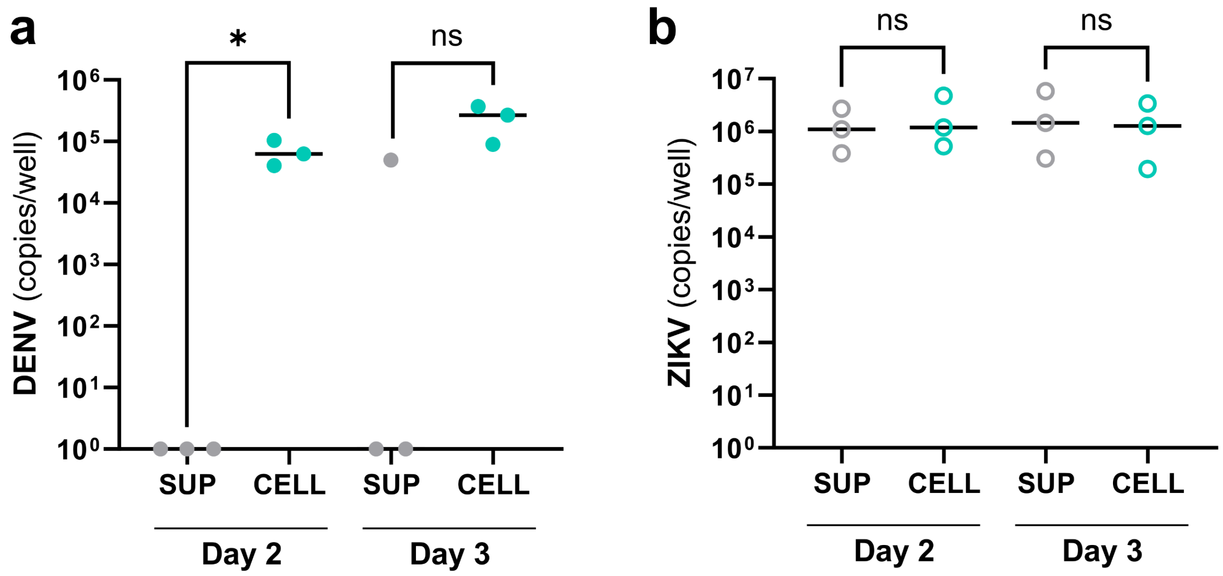

**Figure S3. *In vitro* infections using acellular and cellular hemolymph fractions.** Acellular (SUP) and cellular (CELL) fractions were isolated from the hemolymph of DENV- or ZIKV-infected mosquitoes at 10 days post-infection and then used to challenge C6/36 cells. At 2- and 3-days post-challenge, total RNA was extracted from cells and DENV (**a**) or ZIKV (**b**) RNA titers were determined by qRT-PCR. Each dot represents the viral titer of individual samples with the median displayed by the black line. Data from three independent biological experiments were analyzed using a two-tailed Student's t-test. Asterisks denote significance (*$P$ <0.05). ns, not significant.

**Table S1. Summary of blood meal virus titers for infection experiments.**

| Experiment | Virus | Replicate | GC/ml | FFU/ml |
|---|---|---|---|---|
| CLD Midgut | DENV | 1 | $7.8x10^{9}$ | - |
| CLD Midgut | DENV | 2 | $7.7x10^{9}$ | - |
| CLD Midgut | DENV | 3 | - | - |
| CLD Midgut | ZIKV | 1 | $1.7x10^{9}$ | - |
| CLD Midgut | ZIKV | 2 | $7.1x10^{9}$ | - |
| CLD Midgut | ZIKV | 3 | $9.9x10^{9}$ | - |
| CLD dissemination | DENV | 1 | - | $3.0x10^{7}$ |
| CLD dissemination | DENV | 2 | - | $3.7x10^{8}$ |
| CLD dissemination | DENV | 3 | - | $5.0x10^{8}$ |
| CLD dissemination | ZIKV | 1 | - | $1.3x10^{8}$ |
| CLD dissemination | ZIKV | 2 | - | $1.7x10^{8}$ |

GC/ml = gene copies per ml

FFU/ml = focus forming units per ml

- = not determined

**Table S2. Primers used to assess virus titers in midgut and transfer experiments.**

| Primer | Sequence |
|---|---|
| DENV-2 F | 5’-GCATATTGACGCTGGGARAGAC-3’ |
| DENV-2 R | 5’-TTCTGTGCCTGGAATGATGCTG-3’ |
| DENV-2 Probe | 5’-[6FAM]CAGAGATCCTGCTGTC[BHQ1]-3’ |
| ZIKV F | 5’-TTGTCATGATACTGCTGATTGC-3’ |
| ZIKV R | 5’-CCTTCCACAAAGTCCCTATTGC-3’ |
| ZIKV Probe | 5’-[6FAM]CGGCATACAGCATCAGGTGCATAGGAG[BHQ1]-3’ |

**Table S3. Primers used to assess virus titers in dissemination experiments.**

| Primer | Sequence |
|---|---|
| DENV-2 F | 5'-CATGGCCCTKGTGGCG-3' |
| DENV-2 R | 5'-CCCCATCTYTTCAGTATCCCTG-3' |
| DENV-2 Probe | 5'-[FAM] TCCTTCGTTTCCTAACAATCC [BHQ1]-3' |
| ZIKV 1087 F | 5'-CCGCTGCCCAACACAAG-3' |
| ZIKV 1163c R | 5'-CCACTAACGTTCTTTTGCAGACAT-3' |
| ZIKV 1108 Probe | 5'-[FAM] AGCCTACCTTGACAAGCAGTCAGACACTCAA [BHQ1]-3' |
